# Transient beta activity and connectivity during sustained motor behaviour

**DOI:** 10.1101/2021.03.02.433514

**Authors:** Irene Echeverria-Altuna, Andrew J. Quinn, Nahid Zokaei, Mark W. Woolrich, Anna C. Nobre, Freek van Ede

## Abstract

Neural oscillations are thought to play a central role in orchestrating activity states between distant neural populations. In humans, long-range neural connectivity has been particularly well characterised for 13-30 Hz beta activity which becomes phase coupled between the motor cortex and the contralateral muscle during isometric contraction. Based on this and related observations, beta activity and connectivity have been linked to sustaining stable cognitive and motor states – or the ‘status quo’ – in the brain. Recently, however, beta activity has been shown to be short-lived, as opposed to sustained – though so far this has been reported for regional beta activity in tasks without sustained motor demands. Here, we measured magnetoencephalography (MEG) and electromyography (EMG) in 18 human participants performing an isometric-contraction (gripping) task designed to yield sustained behavioural output. If cortico-muscular beta connectivity is directly responsible for sustaining a stable motor state, then beta activity should be (or become) sustained in this context. In contrast, we found that beta activity and connectivity with the downstream muscle were transient, even when participants engaged in sustained gripping. Moreover, we found that sustained motor requirements did not prolong beta-event duration in comparison to rest. These findings suggest that long-range neural synchronisation may entail short ‘bursts’ of frequency-specific connectivity, even when task demands – and behaviour – are sustained.

**Highlights:**

- Trial-average 13-30 Hz beta activity and connectivity with the muscle appear sustained during stable motor behaviour
- Single-trial beta activity and connectivity are short-lived, even when motor behaviour is sustained
- Sustained task demands do not prolong beta-event duration in comparison to resting state

## Introduction

Frequency-specific patterns of neural activity – often referred to as ‘neural oscillations’ – are ubiquitous in the brain and have been postulated to play a central role in orchestrating communication between remote neural populations (Buzsáki, 2009; Fries, 2005; Siegel et al., 2012; Thut et al., 2012; Varela et al., 2001; Vidaurre et al., 2018). Beta activity (13-30 Hz) is one prominent class of such frequency-specific brain activity (Jenkinson & Brown, 2011; Kilavik et al., 2013) and provides an ideal model system for studying long-range neural communication in humans (Schoffelen et al., 2005). During steady isometric muscle contraction, activity in the primate motor cortex is known to become phase-coupled with that of the contralateral muscle resulting in cortico-muscular coherence (CMC) at the beta frequency (Bourguignon et al., 2017; Conway et al., 1995; Salenius et al., 1997; Schoffelen et al., 2005). These and related findings, typically visualised in trial averages, have led to the proposal that beta activity may play an active role in ‘sustaining’ a stable motor state (Androulidakis et al., 2006; Baker, 2007; Gilbertson et al., 2005; Witham et al., 2011) – and this concept has been proposed to extend to sustaining the ‘status quo’ in the cognitive realm (Engel & Fries, 2010).

In parallel, the sustained rhythmic nature of beta activity is increasingly called into question by a rapidly growing number of reports that demonstrate and quantify that, at the level of single trials, beta activity may not be sustained, but instead occurs in short-lived, burst-like events (Bourguignon et al., 2017; Feingold et al., 2015; Heideman et al., 2020; Jones, 2016; Little et al., 2019; Lundqvist et al., 2016; Quinn et al., 2019; Shin et al., 2017; Tinkhauser et al., 2017; van Ede et al., 2018; see also: Jasper & Penfield, 1949; Murthy & Fetz, 1992, 1996). However, so far, this has been demonstrated and quantified primarily in tasks that did not directly require participants to sustain a measurably steady behavioural output. Instead, beta events have been noted in the resting state (Seedat et al., 2020), during preparatory periods of tasks requiring a single behavioural response (Heideman et al., 2020; Little et al., 2019; Rule et al., 2017; Shin et al., 2017) or during periods following behavioural responses (Feingold et al., 2015; Little et al., 2019). As these contexts may not involve or require a sustained neural process, they may not call for the expression of sustained beta rhythms and instead allow for the manifestation of beta as transient events.

Here, we ask whether motor beta activity is similarly short-lived during an isometric-contraction task designed to yield a sustained behavioural output. If cortico-muscular beta connectivity is directly responsible for sustaining steady motor contraction, then beta activity should be (or become) sustained – also in single trials – in this set-up.

## Results

We measured electrical activity from the brain (using selected magnetoencephalography (MEG) channels over bilateral motor cortices) and the contralateral forearm muscles (using electromyography (EMG) from both forearms), while participants (N = 18) performed a task with instructed periods of bimanual, isometric contraction. Participants held a gripper device in each hand and a prompt instructed them to contract both grippers until they reached the force level indicated on the screen in front of them. They sustained the steady force output (on which they received real-time visual feedback) until the signal to release the grip, 3 s after grip instruction (**Figure 1a**). As shown in the trial-average gripper data from a representative participant (**Figure 1b**; see also **Supplementary Figure 1** for the data from all participants), gripping stabilised approximately 1 second after grip instruction, and was held steady until the instruction to release the grip.

**Figure 1.**
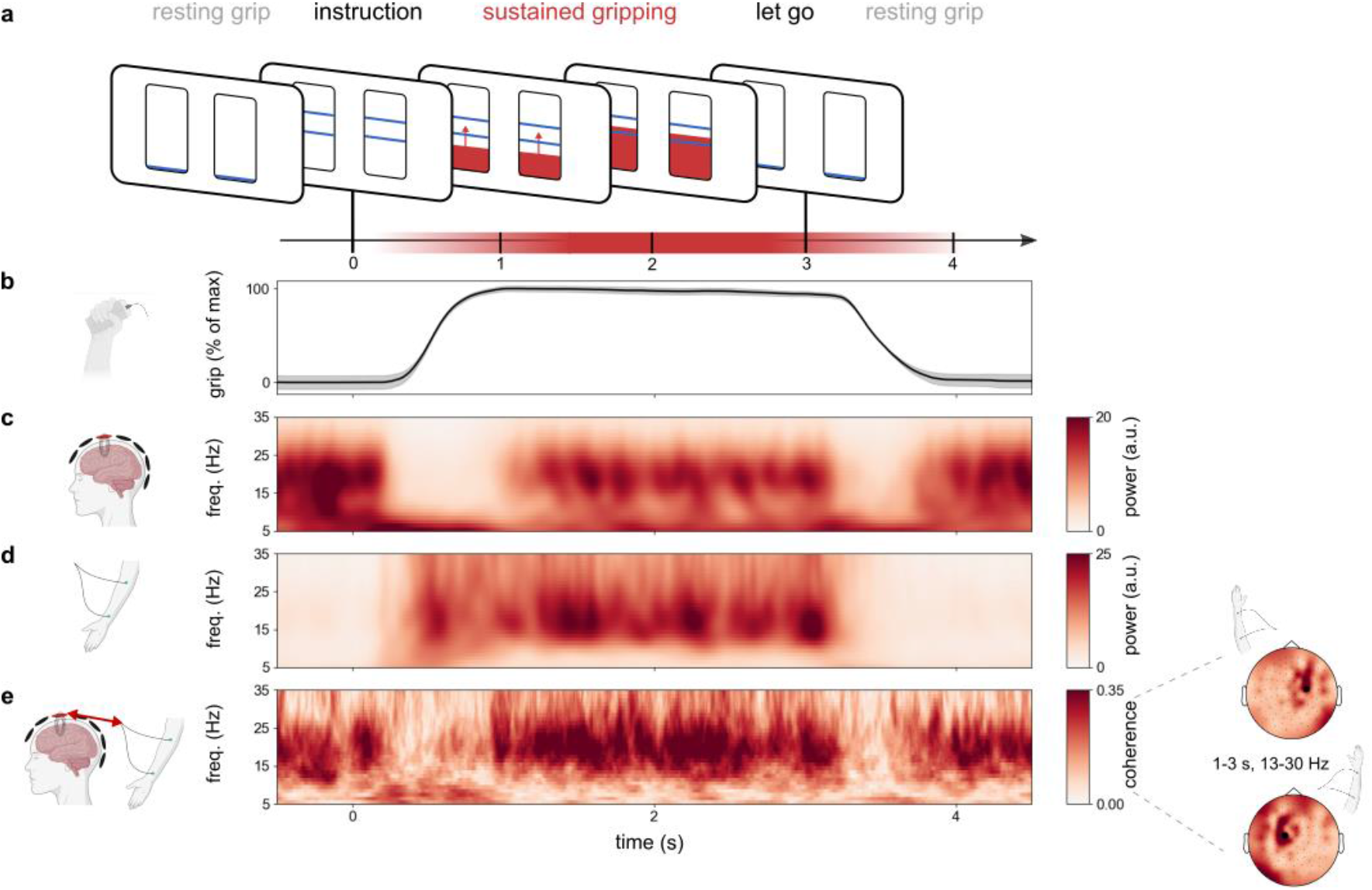
Trial-average beta activity and connectivity appear sustained during sustained motor behaviour (gripping). **a**) Schematic of a single trial. Before each trial, participants held the gripping devices in both hands (resting grip). At time 0, two horizontal lines, indicating the gripping strength, appeared on the bars on the screen, prompting participants to grip. Participants began gripping until they reached a steady grip at the indicated strength, which they sustained for ~2s. The drop of the horizontal lines to the bottom of the bars indicated the end of a trial and the return to resting grip. **b**) Average gripping force across trials and across left and right gripping devices in a representative participant, expressed as a percentage of the maximum force. Shading indicates 95% confidence intervals. **c**) Time-frequency spectrum of trial-average activity in selected motor MEG channels in a representative participant. Selected MEG channels correspond to those with maximal cortico-muscular coherence during sustained gripping (see methods) and are located over the left and right motor cortices, as shown in the topographical distribution in **Figure 1e** (black dots). **d**) Time-frequency spectrum showing trial-average EMG activity across both forearms in a representative participant. **e**) Time-frequency spectrum of cortico-muscular coherence (phase coupling) between the selected MEG channels and the contralateral forearm muscles. Topographies show coherence with the left and right forearms, averaged over the indicated time-frequency window in a representative participant.

### Motor beta activity and connectivity appear sustained during sustained motor behaviour

Similar to previous reports of sensorimotor beta activity during isometric contraction tasks (e.g., Conway et al., 1995; Salenius et al., 1997; Schoffelen et al., 2005), beta power in the brain (**Figure 1c**) and in the muscle (**Figure 1d**) was prominent during the ~2s of stable gripping across trials – from around 1 to 3 seconds after grip instruction. During this period, beta activity in the brain and muscle were also directly related, as revealed by pronounced phase coupling between the motor cortex MEG channel and the contralateral muscle EMG signal at the beta frequency band (cortico-muscular coherence; **Figure 1e** for a representative participant; see **Supplementary Figure 2** for average across all 18 participants). These findings are in line with the putative role of beta in sustaining a steady output or ‘status quo’.

**Figure 2.**
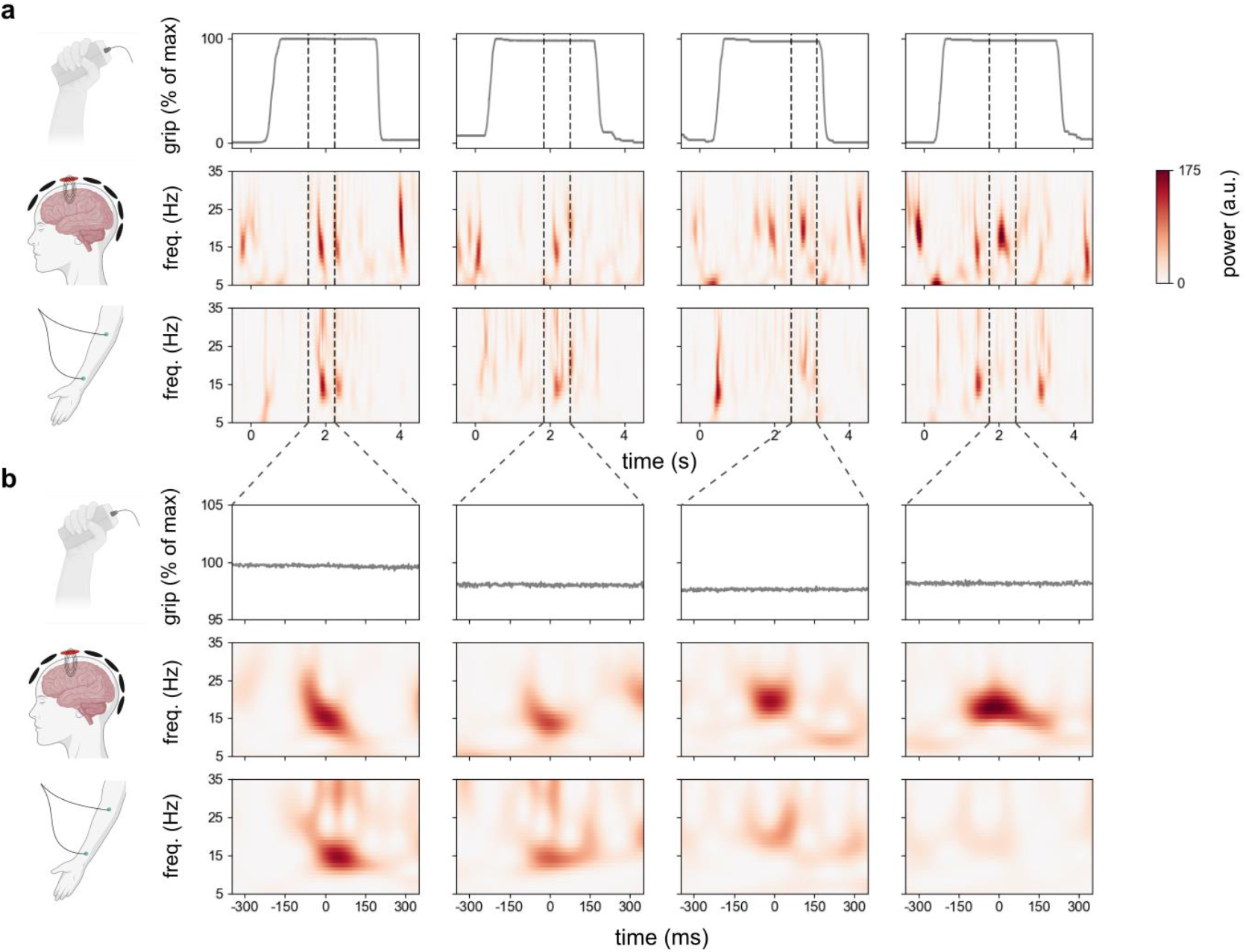
Beta activity is transient in single trials, despite sustained motor output. **a**) Gripper traces together with time-frequency spectra in brain and muscle in four example trials from the same participant whose trial-average data is shown in **Figure 1**. **b**) Zoomed in view of the data in **a**, aligned to the occurrence of a beta event in the single trials above. Percentage of maximal grip output was defined relative to the whole epoch.

In contrast to the period of steady contraction, beta activity in the brain, and its connectivity with the contralateral muscle, were relatively attenuated when participants’ grip force changed – either from the resting grip to the instructed level, or from the instructed level back to the resting grip – similar to previous reports (Salmelin & Hari, 1994).

### Beta activity is transient in single trials, despite sustained motor output

Trial averaging can lead to seemingly sustained activity even when the single-trial activity is short-lived (Jones, 2016; Lundqvist et al., 2016; Stokes & Spaak, 2016). We therefore next turned to the patterns of beta activity at the level of individual trials (**Figure 2**).

Participants’ grip-force traces were sustained throughout the gripping period in the majority of trials (see **Figure 2a** for some representative trials, see also **Supplementary Figure 3**). We also found prominent periods of beta activity at the level of single trials, in both the brain (**Figure 2a**, middle row) and the muscle (**Figure 2a**, lower row). Critically, however, unlike the measured grip, beta activity appeared highly transient at the single-trial level in both the brain and the muscle. In fact, after exhaustive visual inspection of single trials, we found no trial with a clearly sustained period of pronounced beta activity (**Supplementary Figure 3**).

**Figure 3.**
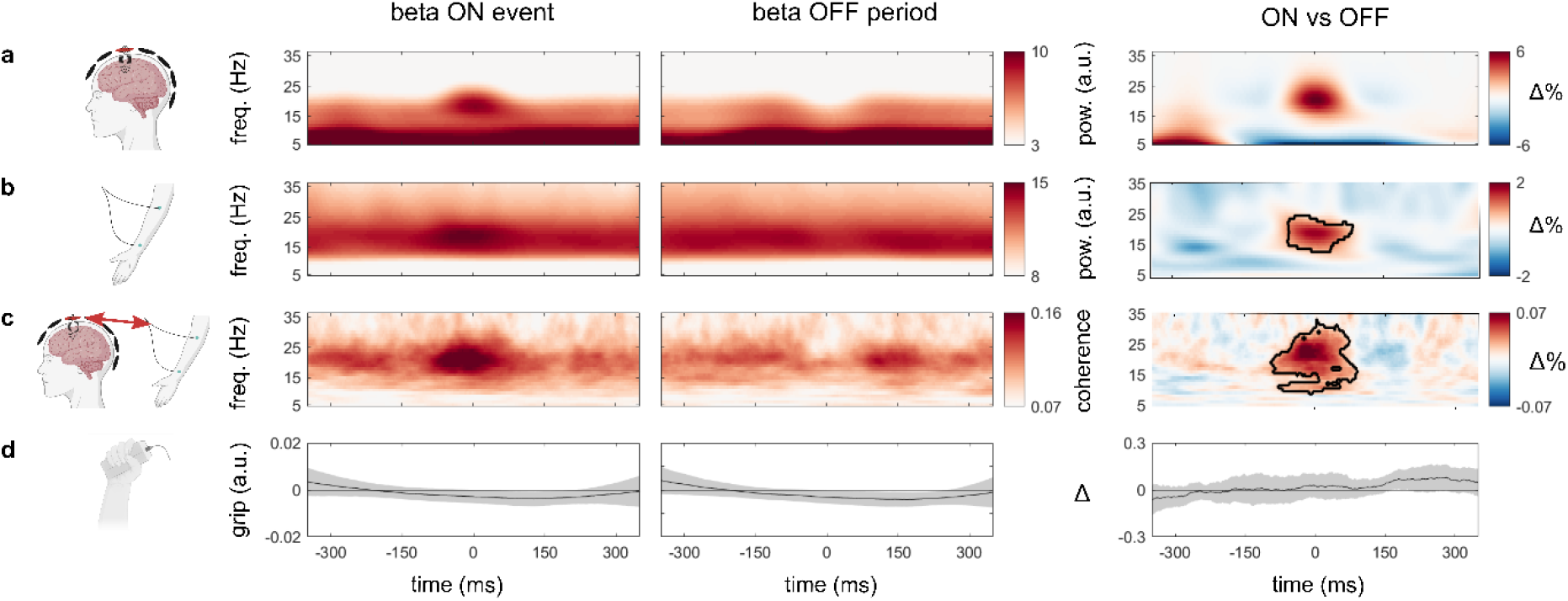
Data aligned to beta ON and OFF periods in the brain reveal transient beta in brain, muscle, and connectivity despite sustained motor output. Beta ON events and OFF periods were identified using Empirical Mode Decomposition (EMD) in the two MEG motor channels during the ~1-3s period of sustained gripping. After ON and OFF period detection, data were aligned and averaged across all trials and all participants (N=18). Columns from left to right represent our four signals of interest (**a-d**) aligned to the central point of a beta ON event (left), a beta OFF period (middle) and the difference between them (right). **a**) MEG time-frequency spectrum, **b**) EMG time-frequency spectrum, **c**) CMC time-frequency spectrum and **d**) gripper signal. Shading indicates 95% confidence intervals. Black outlines in the right time-frequency maps indicate significant clusters.

**Figure 2b** shows the data aligned to prominent individual ‘beta events’ from **Figure 2a**. These data revealed several important points, which we further describe and quantify below. First, periods of high beta power were often transient, lasting a few hundred milliseconds at most. Second, during contraction, periods of high beta power in the brain (middle row) were often accompanied by a similarly short-lasting period of beta activity in the downstream contralateral muscle (bottom row).

Finally, transient beta activity appeared to occur despite sustained motor output, making it unlikely that the observed transient nature of beta was a direct consequence of (undesired) transient motor behaviour during our task (e.g. corrections in grip).

To quantify this pattern across all of our data, we identified periods of high beta power (‘ON’ events) in the single-trial MEG data from the pre-defined channels (that were sensitive to motor activity in the brain), and accordingly aligned the wavelet-transformed MEG, EMG and CMC data, as well as the grip output (**Figure 3**).

For event detection, we used Empirical Mode Decomposition (EMD), a data-driven signal decomposition technique, which has the benefit of preserving high temporal resolution when exploring frequency-specific patterns of brain activity (Huang et al., 1998; see Methods and **Supplementary Figure 4** for details). EMD allowed us not only to capture beta activity in the MEG signal with high temporal resolution, but also to successfully isolate it from other frequency bands (**Supplementary Figure 5**). We note, however, that the advantages of using EMD for beta-event detection were not a pre-requisite for obtaining our central results (as presented in **Figure 3**). Similar results could be obtained when using a simple median split on beta frequency power for beta-event detection (**Supplementary Figure 6**).

**Figure 4.**
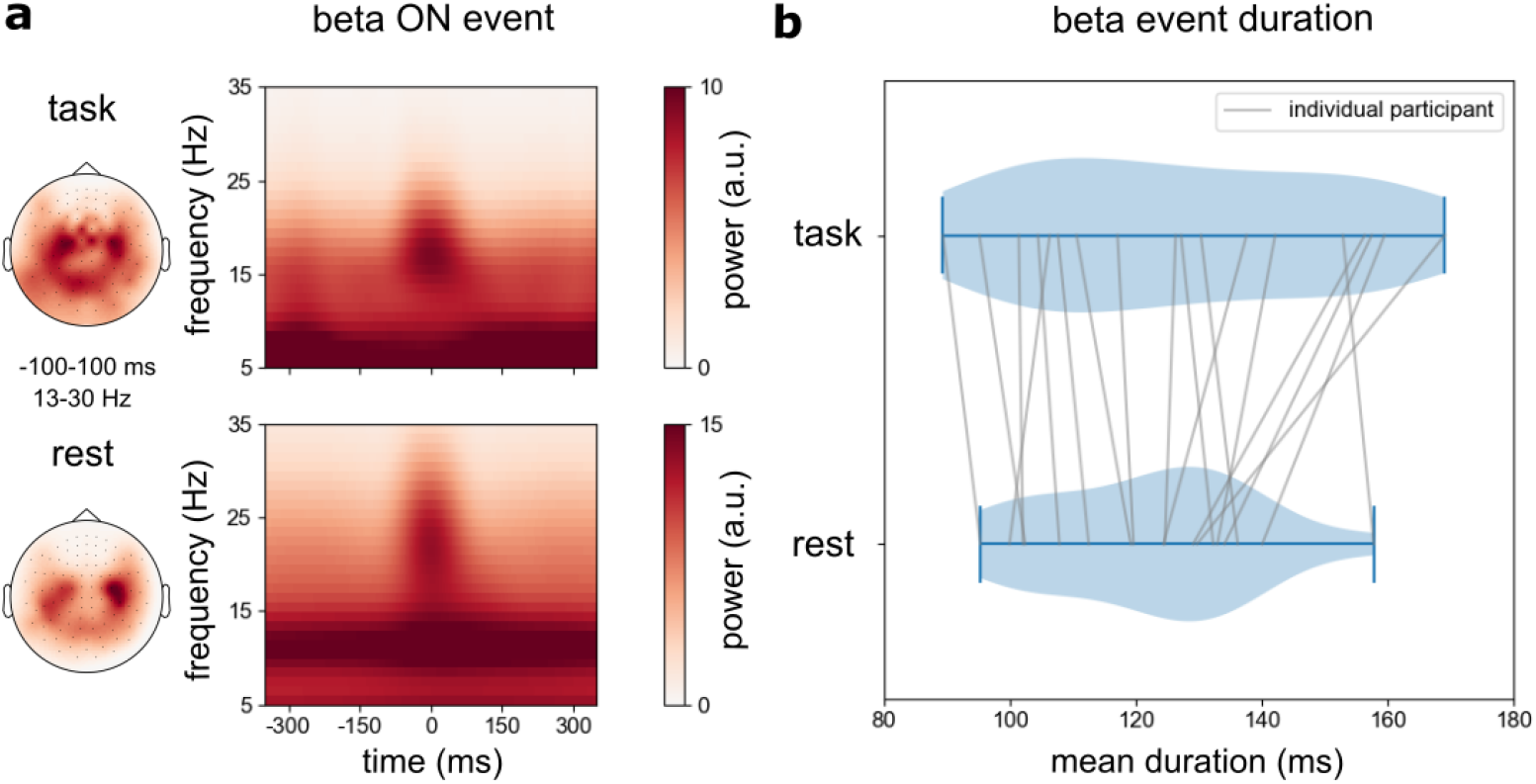
Comparison of the mean duration of beta events during sustained gripping and rest. **a**) MEG time-frequency spectra aligned to all beta ON periods during sustained gripping (task) and during resting state. Topographies show beta power (13-30 Hz) at the time of the detected ON events during gripping and rest. **b**) Violin plot showing the distribution of mean beta event durations during gripping (top) and resting state (bottom). Grey lines represent individual participants. A paired-sample t-test revealed no significant differences in mean burst duration between periods of sustained motor contraction and periods of rest (t_17_ = 1.52, p = 0.15, d = 0.36).

To focus on the sustained gripping period of interest, we only sampled beta events that occurred during the steady-gripping period (as depicted in **Supplementary Figure 1**). For data alignment, we only considered ON events that lasted at least 50 ms (the duration of a single beta cycle at 20 Hz). On average we detected 464.9 ± 114 [M±SD] usable ON events per participant. For comparison, we also identified ‘OFF’ periods, defined as periods in-between the ON events. To compare neural activity surrounding all ON and OFF periods, we aligned our wavelet-transformed data to the centre time-point of all ON and OFF periods.

When averaging across all the identified beta ON events (**Figure 3,** left column) and beta OFF periods (**Figure 3,** middle column) across all trials and all participants, we confirmed that the identified beta ON events were short-lived. This was particularly clear when comparing ON vs OFF periods (**Figure 3**, right column). In line with our single-trial observations (**Figure 2**), we further confirmed that beta ON events in the brain (**Figure 3a**) were accompanied by a correspondingly short-lasting increase in beta activity in the muscle (**Figure 3b**), and by a similarly time-limited pattern of beta connectivity between brain and muscle (**Figure 3c**). The observed patterns did not critically depend on our use of EMD to define ON and OFF periods, as similar results were obtained when defining ON and OFF periods based on a simple median split of 13-30 Hz power time-courses (**Supplementary Figure 6**).

Cluster-based permutation analyses (Maris & Oostenveld, 2007) confirmed significant, transient differences between ON vs OFF periods (as defined in the brain) in beta activity in the muscle (**Figure 3b** third column; cluster p = 0.023) and in the connectivity between the brain and the muscle (**Figure 3c** third column; cluster p = 0.0002). These clusters appeared in frequencies within the beta band and with durations corresponding to those of the beta ON events identified in the brain. Importantly, the significant clusters on the muscle and the brain-muscle connectivity were found despite the fact that these data were aligned to the beta ON events that we identified exclusively on the brain activity (to avoid double dipping, we did not statistically evaluate the ON-vs-OFF effect in the MEG data).

The physiological difference measured in the brain and muscle, as well as in their connectivity, during beta ON periods did not appear to be driven by transient changes in grip output by the participants. Gripping-force differences between beta ON and OFF periods showed no clear modulation around the time of the identified ON/OFF periods and were never significantly different from zero (no clusters found; all uncorrected p >= 0.22). Together, these findings suggest that motor-cortex beta-frequency activity, and its connectivity with the contralateral forearm muscle, is transient, even when participants are sustaining a steady grip.

### Beta events are similar in duration during sustained motor output and rest

While beta ON events were short-lived in our task, it was possible that they were *more* sustained (i.e., longer) than the transient beta events that have previously been reported in the absence of sustained behavioural output. To address this question, we used the same analysis pipeline to identify beta ON events in MEG recordings of the same participants during a separate resting-state recording.

As shown in **Figure 4a**, the beta events in the brain (as identified by our analysis) were similar in duration, as well as topography, during the sustained contraction task (top) and during a resting-state recording (bottom). When directly comparing the mean duration of all identified beta ON events per participant (**Figure 2b**), no significant changes in beta event duration could be identified (t = 1.52, p = 0.15).

## Discussion

Our data confirm previous findings that beta activity varies with overall motor-task requirements (Baker, 2007; Gilbertson et al., 2005; Jenkinson & Brown, 2011; Kilavik et al., 2013; Kilner et al., 2000; Little et al., 2019). At the same time, they also reveal an important dissociation in duration. Beta activity, and its connectivity with the downstream muscle, are manifest as transient events even when participants engage in sustained motor behaviour. Reinforcing the temporal dissociation, we found no obvious correspondence between the occurrence of a beta event – in the brain, the muscle, and their connectivity – and a change in overall force output surrounding the identified beta event. That is, while beta events identified in the brain had a clear correspondence in the pattern of muscle activity (see also Bourguignon et al., 2017; Novembre et al., 2019; Seedat et al., 2020; Tomassini et al., 2020), grip output was similarly sustained in periods with and without clear beta events.

This lack of a direct mirroring between beta activity and behaviour (for other examples see also Murthy & Fetz, 1996; Rule et al., 2017; van Ede et al., 2015) is unlikely to result from lack of sensitivity: we were able to identify and aggregate more than 8,000 ON events, and the ON events were clearly distinguishable from OFF periods (in the brain, muscle, and connectivity). Furthermore, the gripper signals did show robust and pronounced changes in grip strength during the instructed grip periods, at the single-trial level.

A widely held theoretical position is that frequency-specific patterns of brain activity play a role in coordinating communication between distant neural populations (Buzsáki, 2009; Fries, 2005; Siegel et al., 2012; Varela et al., 2001). If true, our data reveal how such long-range synchronisation may entail short ‘bursts’ of frequency-specific connectivity that may last only a few cycles (see also Baker et al., 2014; Bourguignon et al., 2017; Vidaurre et al., 2018; Seedat et al., 2020). Our results further emphasize that this may be the case even when task demands call for sustained behavioural output – and that sustained task demands do not prolong beta-event duration in comparison to rest. In our view, however, this need not imply that beta activity plays no role in sustaining neural communication and behaviour. Indeed, short ‘bursts’ of beta activity and connectivity may contribute to sustained activity indirectly, such as by probing the state of the periphery (Baker, 2007; Witham et al., 2011) and/or triggering a separate neural process that then implements sustained behaviour.

At least one prior study has reported that beta activity in motor cortex may ‘emerge’ transiently in LFP measurements, even when neural firing rates in the same areas are sustained at the beta frequency (Rule et al., 2017). Their findings highlight that LFP signals provide complementary information to neural spiking. Accordingly, transient activity observed in our MEG measurements need not imply that there is no sustained beta rhythmicity in other aspects of neural activity to which our measurements were simply insensitive. Perhaps, then, the most urgent question prompted by the current work pertains to how macroscopic beta activity and connectivity relate to other neural processes at the more granular level, and how these contribute to sustaining neural interactions and behaviour. At a minimum, our data reveal that the links between these respective processes may be far less straightforward than commonly assumed.

Here, we focused on beta activity in the human sensorimotor system during a simple sustained motor task. Related work in non-human primates has shown how beta (as well as gamma) activity in prefrontal cortex during working memory maintenance may be similarly transient (Lundqvist et al., 2016). This is another example of a transient neural activity pattern that occurs despite sustained task demands. However, unlike in our task, sustained demands in working-memory tasks can only be inferred indirectly by assuming that memory maintenance requires sustained activity (an assumption that is increasingly questioned, e.g. Lundqvist et al., 2016; Mongillo et al., 2008; Stokes, 2015). In contrast, by investigating beta activity in the human motor system, we were able to monitor the sustained process of interest (gripping) directly. Doing so, we have revealed that beta activity, as well as connectivity, in the human motor system are transient, even while engaging in measurably sustained behaviour.

## Methods

### Participants

The present study had approval of the Oxfordshire Research Ethics Committee as part of the National Research Ethics Service (Reference number 12/SC/0650). Participants gave informed consent prior to participation and received a monetary compensation upon completion of the study. The data come from a cohort of 18 healthy participants (8 females) of advanced age (M = 69.6 years, SD = 4.66, range = 62-88) who participated as a control group in the context of a larger translational study (Zokaei et al., in revision). All participants except for two were right-handed. Participants spent an average of 16.06 years in education (SD: 2.95). Though the participant sample analysed here is on average older than a typical sample of undergraduate students, the participants understood the task without any trouble and were able to sustain their grip as instructed (**Supplementary Figure 1**).

### Experimental setup

Brain activity was measured using the Elekta NeuroMag 306-channel MEG system at the Oxford Centre for Human Brain Activity (OHBA). Head position inside the scanner was monitored by means of a magnetic Polhemus FastTrack 3D system. Data were acquired with a sampling frequency of 1000 Hz and using a band-pass filter between 0.03 and 300 Hz. Electrocardiography (ECG), electrooculography (EOG) and bilateral electromyography (EMG) were acquired concurrently. EMG electrodes were placed over *flexor digitorum superficialis* (forearm) in a bipolar configuration, with a reference electrode on the lateral epicondyles of each arm (similar to Proudfoot et al., 2018; van Ede & Maris, 2013).

Participants were presented with visual stimuli on a 58×46 cm screen which was 120 cm away and which had a spatial resolution of 1280×1024 pixels. The task was programmed and implemented using Presentation (Neurobehavioral Systems, Inc., Berkeley, CA, www.neurobs.com). Participants’ responses on each trial were recorded using a bimanual fibre-optic gripping device that measured gripping strength and which was compatible with the MEG scanner (Grip Force, Force Fibre Optic Response Pad, Current Designs, USA).

### Stimuli and experimental procedure

Participants were asked to hold the gripping device in both hands for the duration of the experiment. The task consisted of 12 blocks (each separated by a 20-s rest period) with each 10 trials of bimanual gripping: a total of 120 trials per participant. Each gripping trial lasted 3 s and the whole task lasted around 10 minutes. At all points during the task, the screen showed two bars, each associated with one of the hand-held gripping devices (**Figure 1a**). At time 0 of each trial, one of two potential grip strengths (low or moderate) was indicated by two perpendicular lines on the bars, with lower or higher positions representing low and moderate grip strengths. Indicated grip strength was always the same on both bars (i.e. for both hands). After appearance of the two horizontal lines, participants were asked to grip the grippers and sustain their grip until the horizontal lines dropped to the bottom of the bars, at which point they were to release grip (while continuing to hold the gripper devices in their hands). The horizontal lines that indicated the instructed grip strength were on the screen for 3 s. On average, allowing for rise time, participants’ grips were steady at the indicated strength for approximately of 2 s (**Figure 1b**; **Supplementary Figure 1**). During sustained gripping, participants received real-time visual feedback of the strength with which they were gripping, with the aim of ensuring a stable, steady grip strength across the trial. For the purposes of the present study, low and moderate grip strength conditions were collapsed and analysed together. We confirmed that the critical patterns reported in the current study were similar in both grip-strength conditions.

Earlier in the session, the same participants completed a 10-minute resting-state MEG recoring with their eyes open. The same preprocessing and analysis pipeline was applied to the MEG signals recorded during the gripper task and during resting state.

### Data pre-processing

Noisy MEG channels were detected and corrected using MaxFilter version 2.2.15 software (Elekta Neuromag). Other pre-processing steps were performed using FieldTrip toolbox (Oostenveld et al 2011). Artifacts related to eye movements, blinks, and heartbeat were removed using independent-component analysis (ICA) and visual inspection of the detected components. An average of 1.3 components [SD: 1.1] were removed per participant.

We focused our analyses on planar gradiometer MEG channels. Pairs of gradiometers (two per position) were combined in the time domain using the ‘svd’ method in Fieldtrip. To focus our analysis on the temporal dynamics of beta activity in those MEG channels that best captured activity in the human cortico-muscular pathways, we first computed the average corticomuscular coherence (CMC; as in Schoffelen et al., 2005; Salenius et al., 1997) using the data from the full predefined 1-3 s steady gripping period. CMC was calculated using Fourier-transformed EMG and MEG at each MEG channel for all participants as implemented in Fieldtrip. For further analysis, we chose the two MEG channels (one over left motor cortex and one over right motor cortex) that showed largest CMC in the group average. The sustained vs. transient nature of beta activity was not considered when making this channel selection, as the full gripping period was subjected to our CMC analysis at this initial channel-selection stage. All of the MEG analyses on this study, including those on resting-state recordings, were performed using the same two MEG channels of interest in all participants.

Subsequently, noisy trials were identified and removed from the task recording. Data were cut-out into 6 s epochs beginning 1 s before the start of a trial and ending 5 s after. For each participant, the average grip strength change across trials was plotted and a period of steady gripping was identified by visual inspection (**Supplementary Figure 1**). The mean duration of this period was 2.2 s (SD: 0.2 s). The generalised extreme Studentized deviate test for outliers (*Matlab: isoutlier*) was performed for each trial and for each signal type (EMG, MEG, and gripper) during the selected steady-grip time window. Subsequently, bilateral MEG, EMG, and grip force signals were plotted for each of the identified outlier trials and those deemed noisy or unstable by visual inspection were eliminated from all subsequent analyses. An average of 1.5 trials were eliminated per participant (SD: 1.29, range: 0-4).

### Spectral analyses

MEG and EMG time-series were converted into the frequency domain by means of convolving the signals with a complex 5-cycle Morlet wavelet. This was done from 5 to 40 Hz in steps of 1 Hz. Power was calculated as the squared magnitude of the wavelet-convolved data. Time-frequency spectra were subsequently epoched around sustained gripping periods of the task. Time- and frequency resolved CMC was calculated as the consistency in phase-relation between the Morlet-convolved MEG and EMG signals (Schoffelen et al., 2005).

### Detection of ON and OFF periods

Next, we identified periods of high and low beta-frequency (13-30 Hz) activity in the selected MEG channels during the steady gripping period. To identify these periods with high sensitivity while preserving the excellent time resolution of our measurements, we used empirical mode decomposition (EMD) – though we confirmed that similar results could be obtained using a more basic thresholding approach (**Supplementary Figure 6**). EMD is an analysis method developed by Huang et al., (1998, 2003) for the decomposition of nonlinear and nonstationary time series, such as signals arising from brain activity (e.g. MEG). In contrast to conventional Fourier-based time-frequency transforms, EMD allows for higher temporal resolution and it successfully isolates beta from other frequency bands (**Supplementary Figure 5**).

#### Empirical Mode Decomposition

All EMD analyses were performed using Python 3.7 Empirical Mode Decomposition toolbox v0.2.0 (https://emd.readthedocs.io/) together with custom scripts. EMD is based on the empirical (data-driven) identification of a set of intrinsic oscillatory modes in the signal-of-interest and on the decomposition of such signal into a finite set of Intrinsic Mode Functions (IMFs). The algorithm separates the signal *x(t)* into IMFs that fulfil two properties: 1) The number of local maxima in an IMF differs from the number of minima in the same IMF by one or zero, and 2) The mean value of an IMF is zero.

IMFs are isolated through a ‘sifting’ process. In the first sifting step: 1) local minima and maxima of the raw signal are identified, 2) two envelopes connecting all of the maxima and all of the minima of the signal, respectively, are created, 3) the mean of the two envelopes is calculated, 4) this mean is subtracted from the raw signal, 5) if the product of the subtraction itself is *not* an IMF (as defined by the properties above), a new sifting process begins with the product of the subtraction as the new input signal.

To avoid mode mixing (the presence of components at different intrinsic frequencies in a single IMF; Deering & Kaiser, 2005), we used an adaptive version of the sift process similar to the one developed by Huang et al., (2009): masked EMD (mEMD). At each iteration, a masking signal at a specific frequency was summed with the input signal before extrema identification, ensuring that intermittent signals did not result in component splitting. After extrema identification, envelope interpolation and envelope average subtraction from the input signal, the mask was subtracted from the product signal to return the IMF and continue the sifting process. We chose frequency masks at the following frequencies: 125, 62, 31, 16, 8, 4 and 2 Hz (directing modes between the masks, such as between 31 and 16 Hz). To ensure comparability across participants, we imposed an upper bound on the number of IMFs that the sifting process would return; namely, 6. The IMF that best corresponded to what is commonly described as ‘beta’ was the 3^rd^ IMF in all participants (**Supplementary Figure 5**), which was isolated using the 31 and 16 Hz masks (Quinn et al., in revision).

#### Beta cycle detection

Following identification of the beta mode (beta IMF), we identified periods of high and low beta activity. The outstanding temporal resolution of EMD enabled us to do so at the level of single beta cycles. We identified beta cycles in our signal based on the instantaneous phase and instantaneous amplitude of the beta IMF. We defined ‘good’ beta cycles as those: 1) beginning with an instantaneous phase value between 0 and pi/24 and ending within 2pi – pi/24 and 2pi, 2) with a phase differential bigger than 0 (no phase reversals) and 3) instantaneous amplitude above the 50^th^ percentile of the beta IMF’s instantaneous amplitude for each channel, to ensure that only cycles with a sufficiently high amplitude were included. We defined ‘beta ON events’ as successive ‘good’ beta cycles.

#### ON and OFF period identification

We chose chains made of good beta cycles (ON events) that were at least 50 ms long (the estimated duration of a single beta cycle at 20 Hz) and that occurred at least 500 ms away from the beginning or end of the sustained gripping period identified separately for each participant (**Supplementary Figure 1**). For comparison, in the same gripping period, we also marked all moments in between the ON events (OFF periods). We found the central point of these ON and OFF periods and aligned the time- and frequency-resolved data for all ON and OFF periods to their middle point. We also calculated the percentage change between ON and OFF periods’ wavelet-derived time-frequency spectra as: ((ON-OFF) / OFF) * 100. Prior to the comparison of CMC during ON and OFF periods, we randomly subsampled our data to calculate CMC with the same number of ON and OFF periods, provided that CMC is biased by the number of trials.

Although we had good reasons for using EMD (its high temporal resolution and ability to isolate the beta mode from other frequencies), we also confirmed that the results we obtained were not dependent on using EMD. To this end, we also found ON and OFF periods in the more conventional wavelet-convolved time-frequency spectra, and identified ON events as those periods for which beta power (13-30Hz) exceeded the median beta power for at least 50 ms. Highly similar results were observed (**Supplementary Figure 6**).

### Statistical analyses

We focused our statistical analysis on the ON vs OFF comparison described above, separately for the EMG, CMC, and grip output measurements. We refrained from statistically evaluating the ON vs OFF comparison on the MEG data given that we used the MEG data itself to identify the ON events and OFF periods. We used cluster-based permutation testing (Maris & Oostenveld, 2007) as implemented in FieldTrip. This approach circumvents the multiple comparisons problem that stems from statistical evaluation of multi-dimensional data (in our case, data with a time and frequency dimension) by evaluating clusters of neighbouring samples under a single permutation distribution of the largest cluster. We performed 10,000 permutations with a clustering threshold of a univariate comparison at alpha = 0.05. We also compared the mean duration of beta events between steady contraction and rest. For this analysis, we extracted the mean event duration as the time from the beginning to the end of a ‘chain’ of consecutive beta cycles (as defined above) and used a paired samples t-test to compare mean duration of ON events between conditions of sustained contraction and resting state.

## Acknowledgements

This research was funded by a Wellcome Trust PhD Studentship (102170/Z/13/Z) to I.E., Wellcome Trust grants (098369/Z/12/Z, 106183/Z/14/Z, 215573/Z/19/Z) and the New Therapeutics in Alzheimer’s Diseases (NTAD) study supported by UK MRC and the Dementia Platform UK to M.W.W., a Wellcome Trust Senior Investigator Award (104571/Z/14/Z) and a James S. McDonnell Foundation Understanding Human Cognition Collaborative Award (220020448) to A.C.N, an ERC Starting Grant from the European Research Council (MEMTICIPATION, 850636) to F.v.E. and by the NIHR Oxford Health Biomedical Research Centre. The Wellcome Centre for Integrative Neuroimaging is supported by core funding from the Wellcome Trust (203139/Z/16/Z). For the purpose of open access, the author has applied a CC BY public copyright licence to any Author Accepted Manuscript version arising from this submission. Figure schematics were created with BioRender.com.

## Supplementary Figures

**Figure S1.**
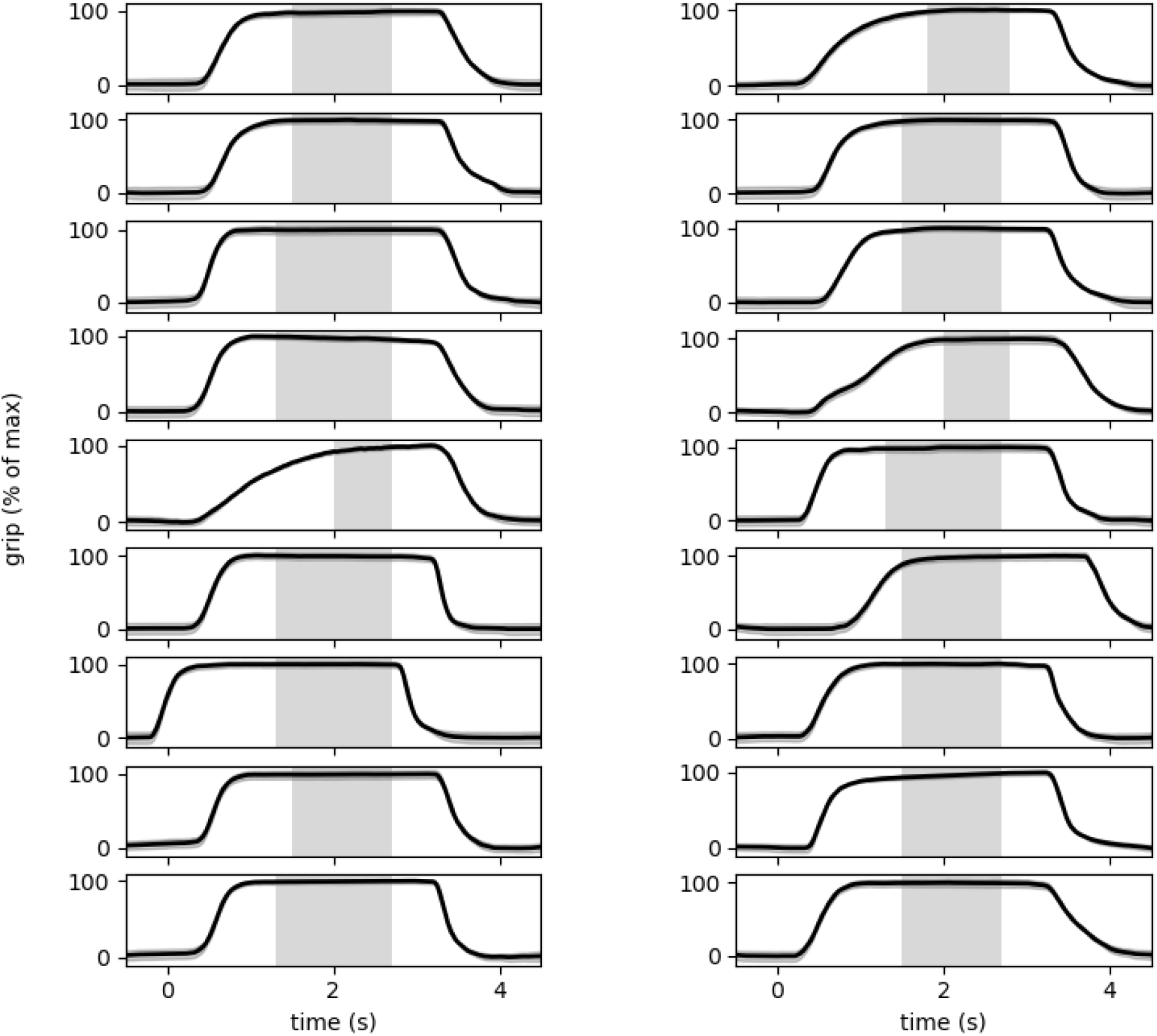
Across-trial average of gripper signal in single participants. The shaded areas around the black lines represent 95% confidence intervals. The shaded rectangular shaded area depicts the central part of the steady gripping period during which beta ON and OFF periods were identified.

**Figure S2.**
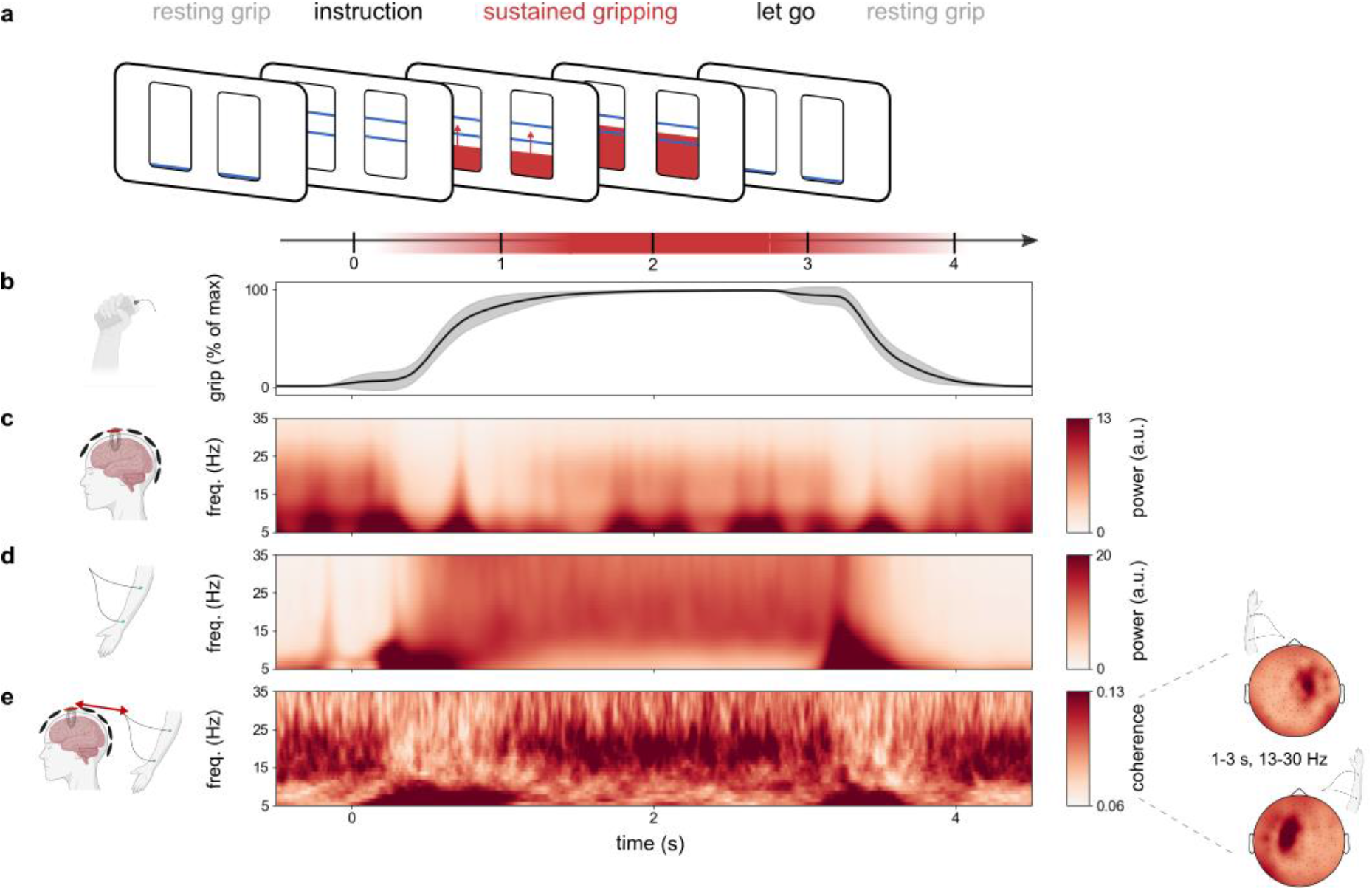
Trial-average beta activity and connectivity appear sustained during sustained motor behaviour (gripping) across participants. **a**) Schematic of a single trial. Before each trial, participants held the gripping devices in both hands (resting grip). At time 0, two horizontal lines, indicating the gripping strength, appeared on the bars on the screen, prompting participants to grip. Participants began gripping until they reached a steady grip at the indicated strength, which they sustained for ~2s. The drop of the horizontal lines to the bottom of the bars indicated the end of a trial and the return to resting grip. **b**) Average gripping force across trials and across left and right gripping devices across participants, expressed as a percentage of the maximum force. Shading indicates 95% confidence intervals. **c**) Time-frequency spectrum of trial-average activity in selected motor MEG channels across participants. Selected MEG channels correspond to those with maximal cortico-muscular coherence during sustained gripping (see methods) and they lie over the left and right motor cortices, as shown in the topographical distribution in **Figure S2e**. **d**) Time-frequency spectrum showing average EMG activity across both forearms across participants. **e**) Time-frequency spectrum of cortico-muscular coherence (phase coupling) between the selected MEG channels and the contralateral forearm muscles. Topographies show coherence with the left and right forearms, averaged over the indicated time-frequency window across participants.

**Figure S3.**
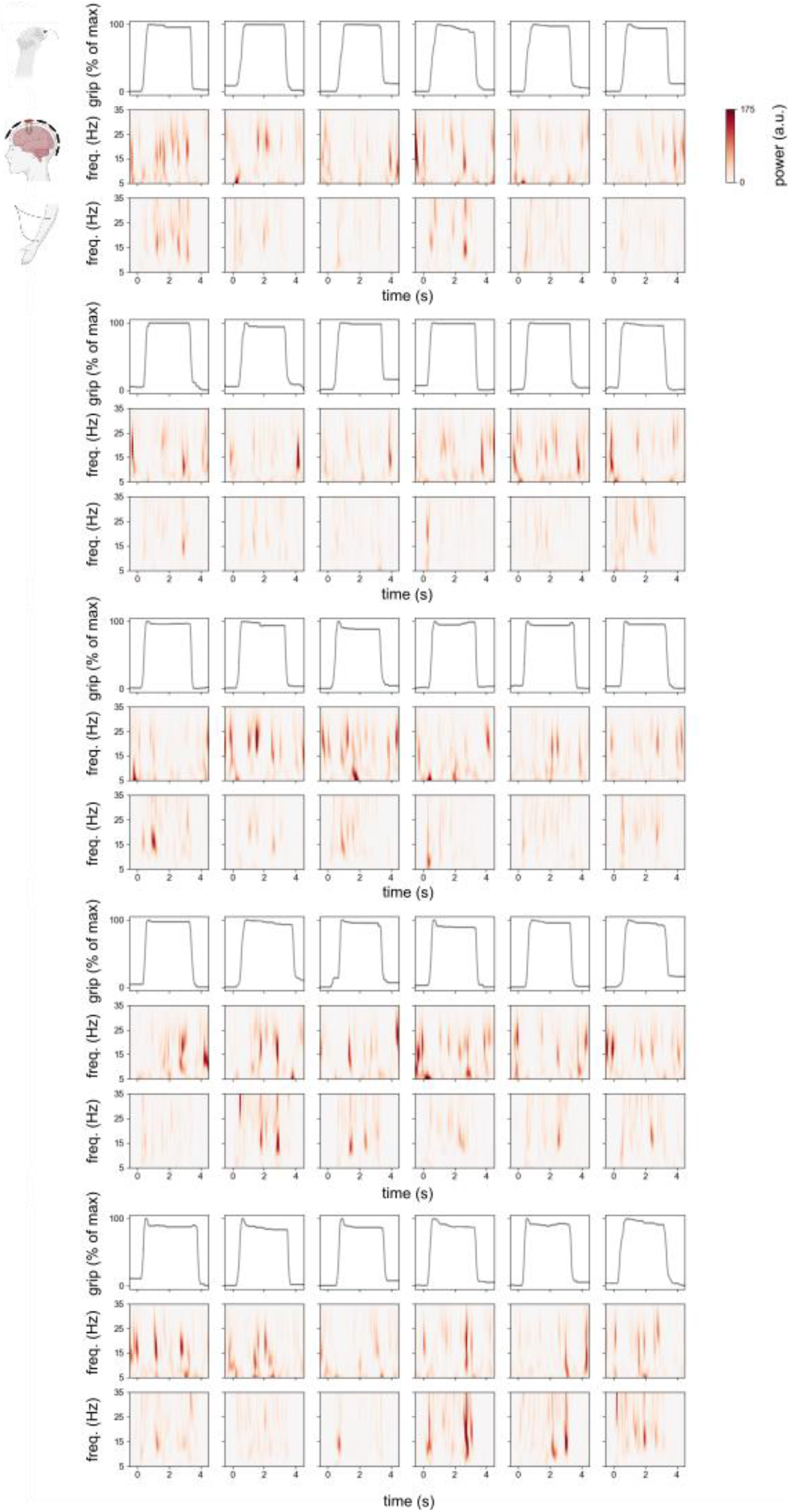
Single trials in the brain, muscle and gripper. Gripper traces together with time-frequency spectra in brain and muscle in 30 example trials from the same participant whose trial-average data is shown in **Figure 2**.

**Figure S4.**
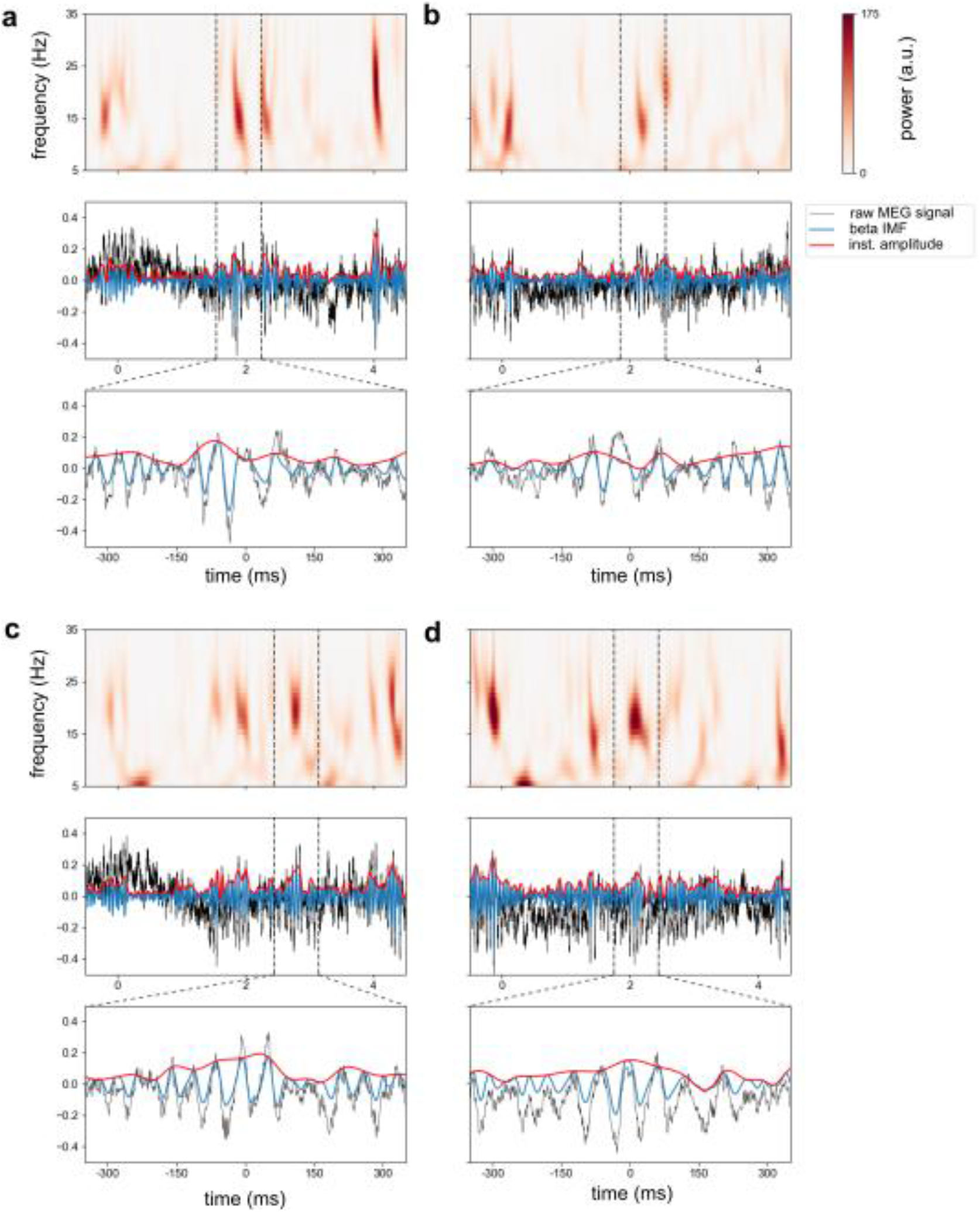
Raw MEG signal, beta IMF and instantaneous amplitude of beta IMF during trials shown in Figure 2. a-d single trials shown in Figure 2. Top: wavelet-convolved time-frequency spectra during the trial. Middle: raw MEG trace (black), beta IMF of the MEG signal (blue), and instantaneous amplitude of beta IMF (red) for the duration of the trial. Dotted lines depict representative individual beta events. Bottom: zoomed in version of the individual beta events.

**Figure S5.**
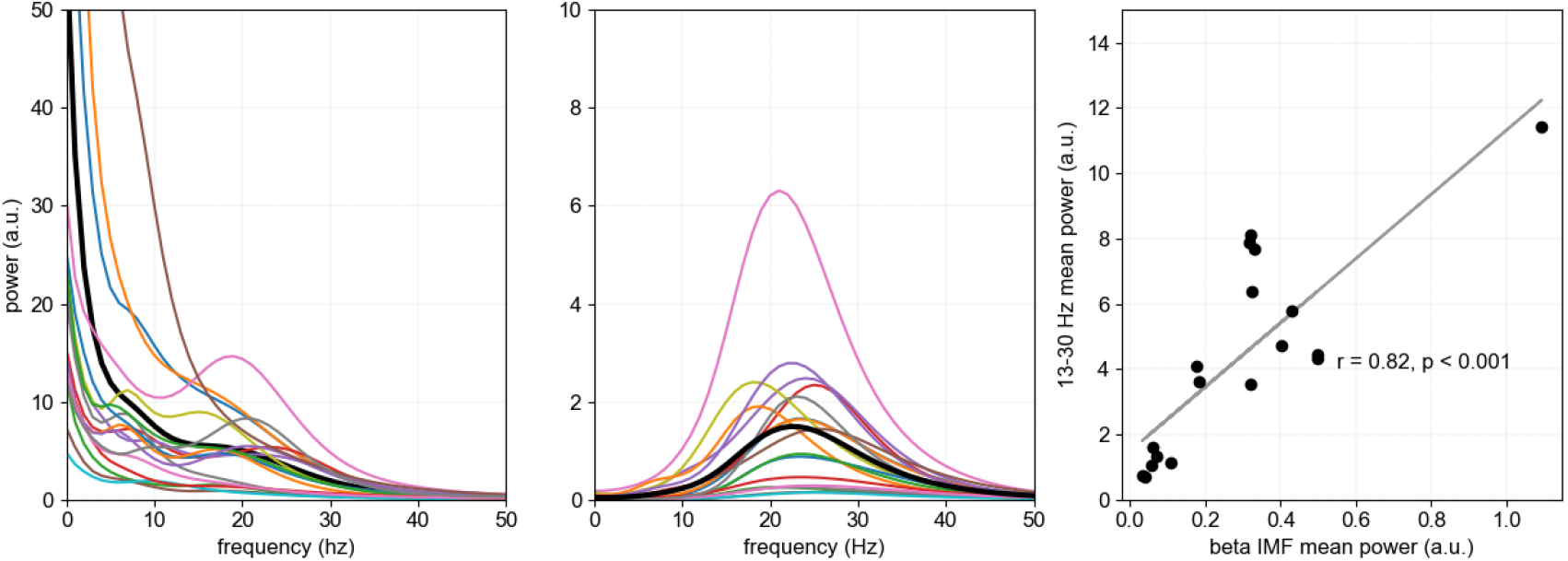
Beta IMF captures beta component and isolates it from other frequency bands. Left: power spectra for each participant (N = 18) and across-participant mean as black line. Middle: power spectra of beta IMF for each participant and average in black. Right: correlation between total power of beta IMF and average power in 13-30Hz frequency range (r = 0.82, p < 0.001).

**Figure S6.**
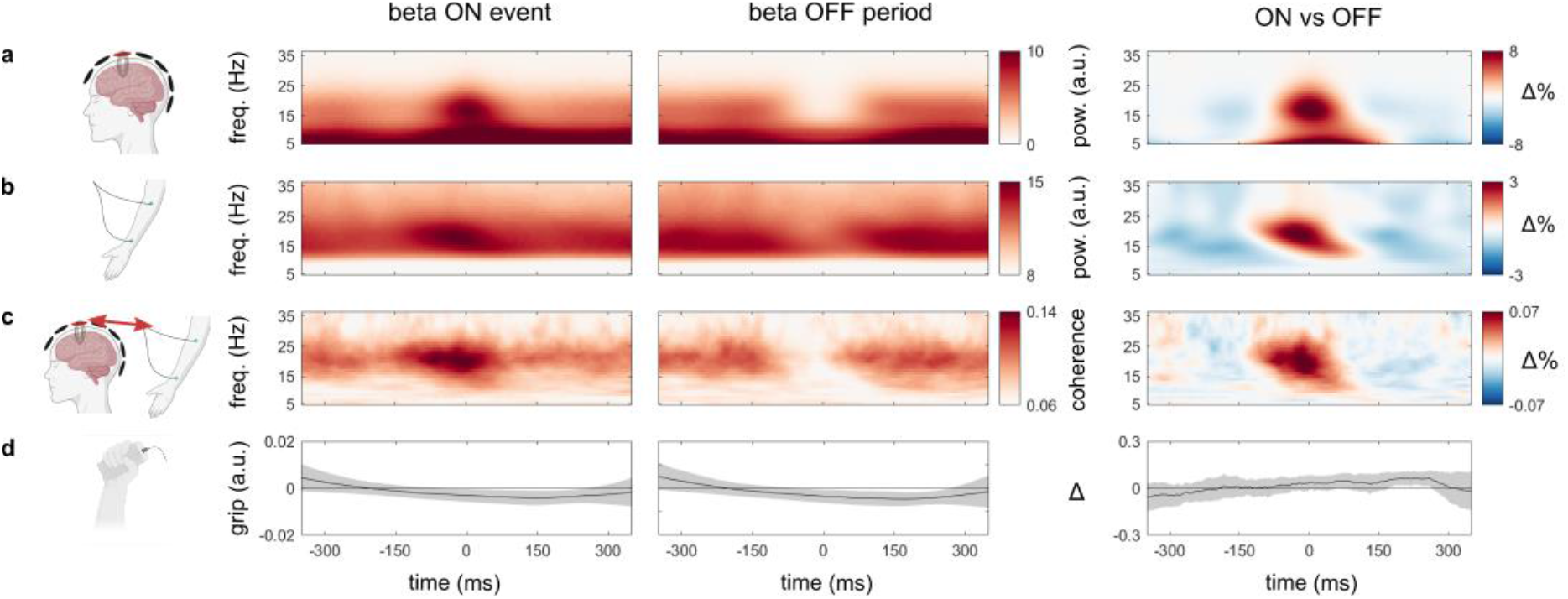
Data aligned to beta ON and OFF periods (identified using a simple median power threshold) in the brain reveal transient beta in brain, muscle, and connectivity despite sustained motor output. Beta ON events were identified as periods of beta power (13-30 Hz) higher than the median beta power per trial in the wavelet-convolved data from the two motor MEG channels during the ~1-3s period of sustained gripping (see methods). Beta OFF periods were periods with beta power below the median. After ON and OFF period detection, data were aligned and averaged across all trials and all participants (N=18). Columns from left to right represent our four signals of interest (**a-d**) aligned to the central point of a beta ON event (left), a beta OFF period (middle) and the difference between them (right). **a**) MEG time-frequency spectrum, **b**) EMG time-frequency spectrum, **c**) CMC time-frequency spectrum and **d**) gripper signal. Shading indicates 95% confidence intervals. Back outlines in the right time-frequency maps indicate significant clusters.

